# Monocyte subpopulations exhibit distinct TNF-dependent aging signatures

**DOI:** 10.1101/2023.11.17.567601

**Authors:** Anna Woo Bentz, Monica Gutierrez, Dessi Loukov, Blair Eckman, Gaurav Gadhvi, Carla Cuda, Dawn M. E. Bowdish, Deborah R. Winter

## Abstract

Monocytes are a key cell type contributing to age-associated inflammation or inflammaging. Since monocytes have the potential to enter circulation and differentiate into macrophages in tissues, they are capable of having a systemic effect on health. Here, we characterize the effect of aging on the transcriptional regulation of monocyte subpopulations in the bone marrow. We find that aging classical (Ly6c high) and non-classical (Ly6c low) monocytes exhibit distinct transcriptional profiles. These were associated with changes to the epigenomic landscape driven by the activity of specific TFs that often had opposing effects in monocyte subpopulations. Next, we determined that the aging signature was diminished in TNF-KO mice indicating that monocytes are altered in a TNF-dependent manner. Finally, we found that a subset of the aging signature was triggered by TNF over-expression. Together, our results implicate key factors driving age-associated changes to the transcriptional regulation of monocyte subpopulations. This study provides a better understanding of the impact of aging on monocytes and identifies targets for future investigation aiming to improve health in aging.

## Introduction

Aging is accompanied by a decline in immune function with aged individuals exhibiting low-grade chronic inflammation^1^. The term “inflamm-aging “was first coined by Franceschi and colleagues to describe the reduced capacity of aged individuals to cope with environmental and molecular stressors^2^. Elevated levels of tissue and circulating pro-inflammatory cytokines are observed in aging even in the absence of an immunological threat^3–5^. Inflammaging significantly contributes to the increased risk of many age-associated morbidities, such as cardiovascular, neurodegenerative, and autoimmune disease.^6–8^ Multiple intrinsic factors are thought to contribute age-associated inflammation including genomic instability, mitochondrial dysfunction, and epigenetic changes ^9–11^. These are compounded by extrinsic signals in the aging microenvironment, such as cytokine signalling and accumulated cellular debris, leading to immunosenescence and increased inflammaging^12, 13^. Understanding the impact of aging on immune cells is critical to formulating therapeutic interventions to halt this feedback loop.

Monocytes and macrophages are key players in the development and propagation inflammaging. In the healthy body, they drive the inflammatory response, activating other cells through the release cytokines, and phagocytosing pathogens, dying cells, and other debris^14^. Monocytes derive from hematopoietic stems cells (HSCs) in the bone marrow and are released into circulation; in inflammatory conditions, they infiltrate into tissue to differentiate into macropahges^15–17^. However, aging leads to mutations and epigenetic changes in HSCs that affect their ability to differentiate^18, 19,20^. The result is that aging hematopoiesis generates a disproportionate number of myeloid cells, including monocytes, in a process called myeloid skewing^21, 22,24^. Thus, monocyte function is altered in aging and their numbers are increased in the circulation of elderly individuals^23–25^. These changes are further transmitted into macrophages: aging macrophage exhibit impaired metabolism, hypersensitivity, and increased senescence ^11, 26, 27^. Macrophages have been implicated in many age-related diseases across a variety of tissues^28, 29^. Therefore, monocytes provide a promising target for therapies aimed at reducing inflammaging.

Monocytes are generally classified as Classical (CM) or Non-classical (NCM) though other subpopulations do exist^16^. These correspond to Ly6c hi vs. Ly6c low in mice and CD14^++^CD16^−^ vs. CD14^+^CD16^+^ in humans ^30, 31^. Murine Ly6c hi monocytes in the bone marrow egress into the blood where they live for a few days or differentiate into Ly6c lo^32, 33^. This process is driven by tightly regulated epigenomic changes that regulate unique expression in CM and NCM^34, 35^. Prior work from our group has shown that Ly6c hi monocytes are preferentially increased in blood, bone marrow, and spleen from aging mice and exhibit distinct surface markers ^36, 37^. Moreover, these effects were diminished in aging mice with depleted tumor necrosis factor (TNF) signalling, such as in TNF knockout mice. We and others have demonstrated that macrophages are capable of responding to signals in their local environment by reprogramming their epigenomic landscape ^38–40^. We further showed that myeloid progenitors and bone marrow derived cells can recapitulate tissue-specific regulatory elements in development or after the niche disruption. Similarly, epigenomic changes accompany monocyte to macrophage differentiation *in vitro* and response to stimulation^41–43^. Thus, we propose that signals, such as TNF, in the aging microenvironment lead to reprogramming of the monocyte epigenomic landscape and transcriptional dysregulation.

Here, we aim to characterize the aging signature of monocyte subpopulations from murine bone marrow. We perform RNA-seq and ATAC-seq on young and aged CM and NCM to profile the transcriptional and epigenomic changes in aging. We identify altered gene expression pathways and chromatin accessibility of cis-regulatory regions to implicate key factors in monocyte aging. We also compare the results from wild-type (WT) monocytes with those isolated from TNF-KO mice to assess TNF dependency. Finally, we assess whether the accessibility age-associated regions are also affected by TNF over-expression. The results of this study provide potential targets to explore for improved health in aging.

## Results

### Classical and Non-classical Monocytes Exhibit Distinct Transcriptional Signatures of Aging

In order to better understand monocyte differentiation by subpopulation, we performed RNA-seq on bone marrow monocytes from young and old mice. We used flow cytometry to sort classical (CM) and non-classical (NCM) monocytes based on surface marker expression of Ly6c (**Supp Figure 1A**). Not surprisingly, the differences between subpopulations represented the largest variability in the transcriptional profiles between samples (**Supp Figure 1B-D**) but the age of mice also drove significant differences in gene expression (**Supp Figure 1E**). We identified 1096 and 562 differentially expressed genes (DEGs) between young and old mice in CM and NCM, respectively (**Figure 1A-C**). Genes with decreased expression in aging CMs included those associated with activation of immune response (Ifi206, Orgm2, Mnda) and defense response to virus (CxCl10, Stat1, Isg15); while those that were increased in expression were associated with translation (Eif3b, Rps19l; Ndufa7) and other metabolic processes (**Figure 1D**). On the other hand, the only GO process significantly associated with decreased expression in aging NCMs was “positive regulation of maintenance of sister chromatid cohesion” (*Ankrd32, Sgol2, H2afy*). Genes with increased expression in aging NCMs included those in the TNF and IL6 pathways (*Cyba, Bcl3, Fcer1g, Nfkbil1*). Only a small proportion of DEGs overlapped in the same direction between NCM and CM although there were similar trends (**Figure 1E**; **Supp Figure 1F**). These results indicate that the transcriptional regulation of monocyte subpopulations is distinctively impacted by aging.

**Figure 1.**
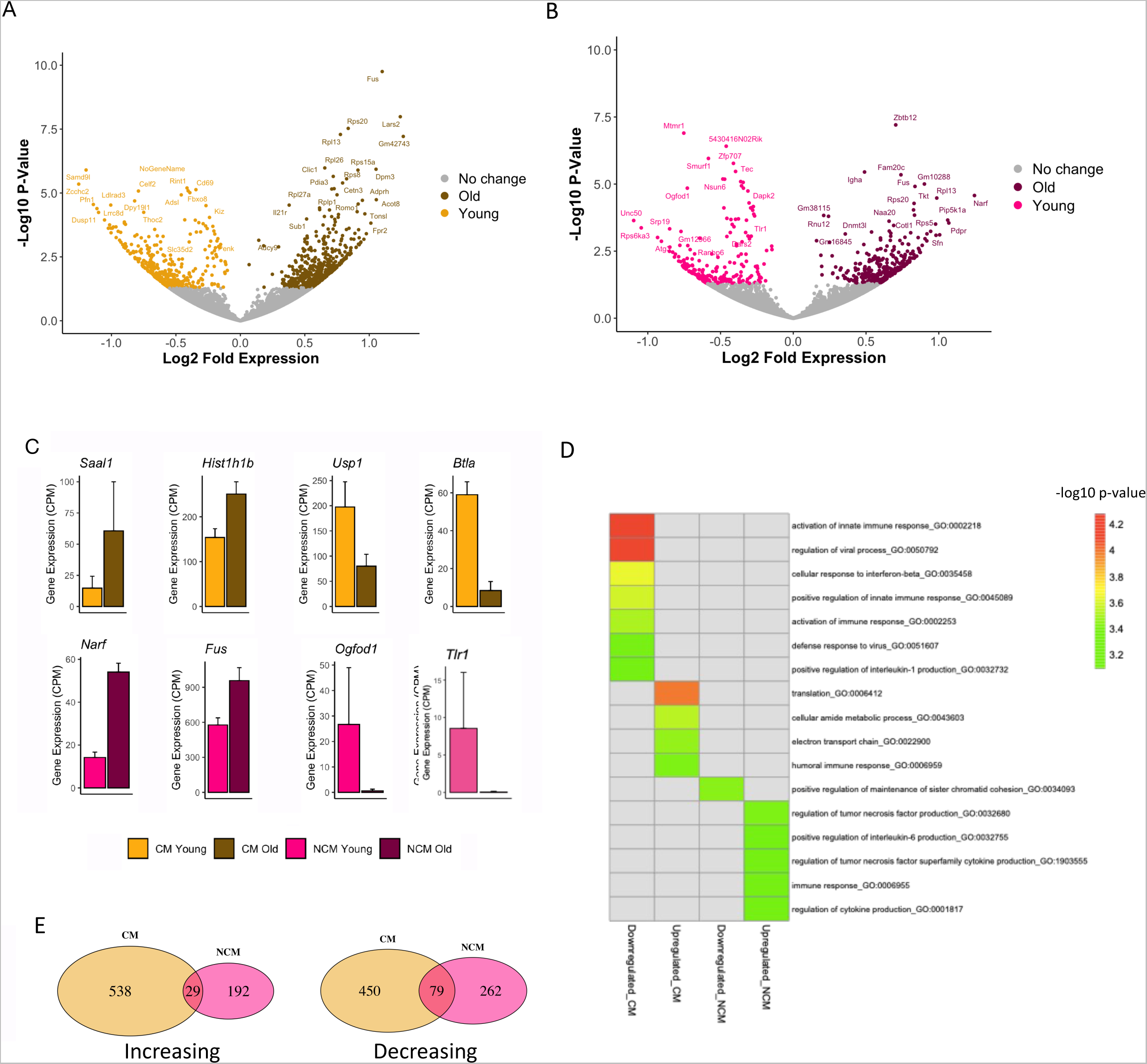
Ly6CHigh and Ly6CLow monocytes display unique age-associated transcriptional profiles. A. Volcano plot of gene expression changes between young and old classical (CM) monocytes. B. Volcano plot of gene expression changes between young and old nonclassical (NCM) monocytes. C. Representative examples of differentially expressed genes (DEGs) in aging from either CM or NCM (CPM counts per million). Error bars represent standard deviation (S.D.) D. Heatmap of gene ontology (GO) analysis results on DEGs in aging from CM (A) and NCM (B). E. Venn diagrams of overlap between aging DEGs with increased (left) or decreased (right) expression in CM and NCM.

### Widespread Epigenomic Changes in Aging Monocytes Linked to Transcription Factor Activity

Epigenetics is one of the hallmarks of aging^44^ and prior studies have shown that cis regulatory elements found in open chromatin regions are critical to monocyte specification^34, 35^. We performed ATAC-seq on monocyte subpopulations from the same young and old mice to profile changes in the epigenomic landscape with aging (**Figure 2A**). We defined 6269 and 2668 regions as part of the aging signature of differential chromatin accessibility in CM and NCM, respectively (**Figure 2B**). Next, we used LOLA to determine whether these differential regions were enriched for experimental binding sites of transcription factors and chromatin regulators. Regions with increased accessibility in aging CM significantly overlapped binding sites of CEBP and AP1 (ATF4, JunB) in macrophages; while those with decreased accessibility overlapped with CTCF and cohesin associated proteins (**Supp Figure 2A**). Interestingly, it was regions that increased in accessibility in aging NCM that overlapped CTCF and cohesin proteins as well as AP-1 family (ATF3). The regions decreased in accessibility in aging NCM overlapped with binding sites associated with polycomb repression (Suz12, Ezh2, Ring1B) in stem cells. Taken together, we observe systemic changes in the chromatin profiles of both CM and NCM.

**Figure 2.**
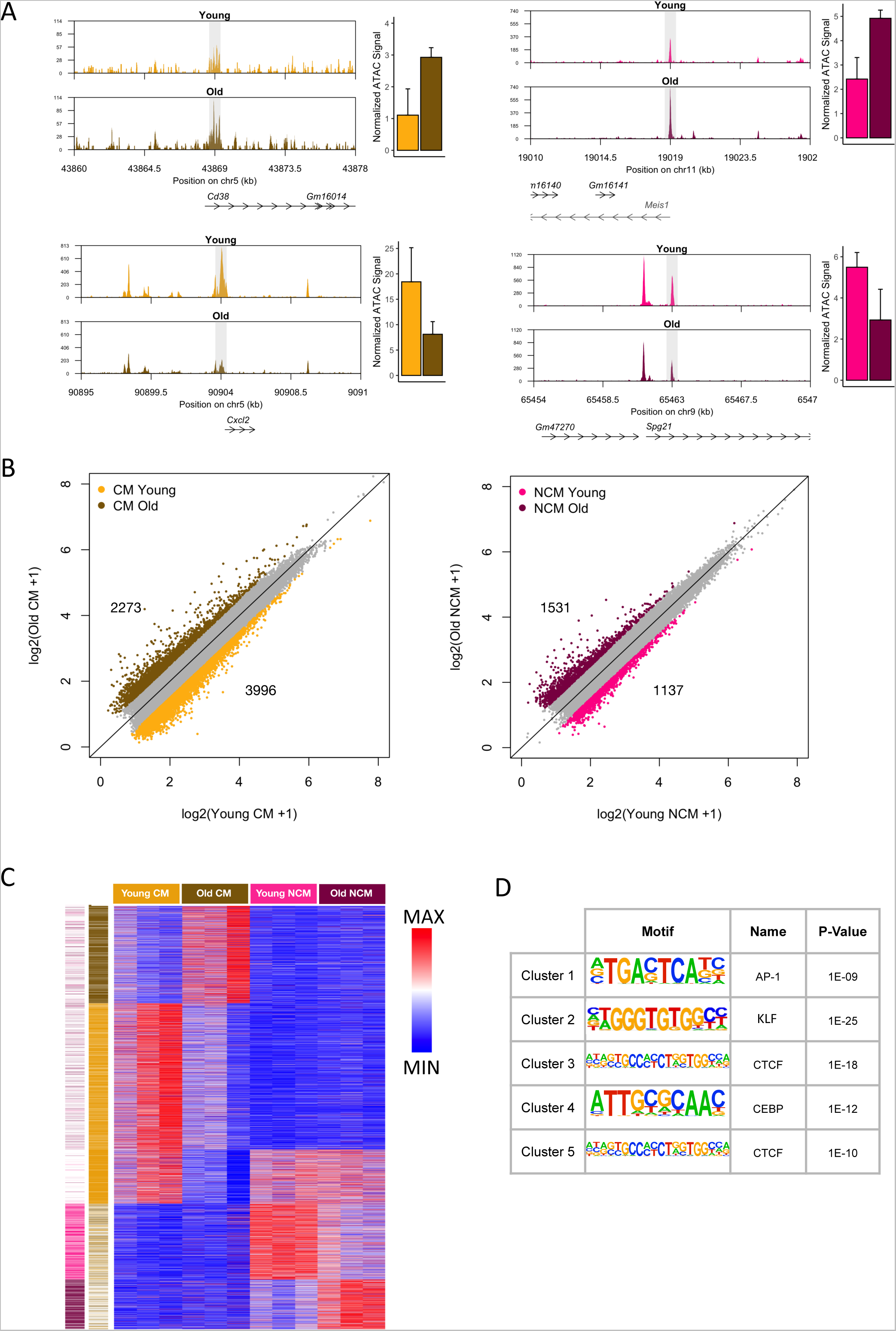
Age dependent chromatin accessibility changes in Ly6CHigh and Ly6CLow monocytes. A. Genome tracks of chromatin accessibility show representative regions that increase (top) and decrease (bottom) in chromatin accessibility with aging in CM (left) and NCM (right). Data in genome tracks represent the average of 3 replicates in RPKM. Mean normalized ATAC signal for highlighted regions are shown to the right. B. Scatter plots of the average ATAC-seq signal in log2 space for young and old CM (left) and NCM (right). Regions with a difference greater than 0.5 are colored and represent the aging signature. C. K-means clustering (k = 5) of chromatin accessibility changes in 8686 regions from the combined CM and NCM aging signature. D. Motif families that are significantly enriched in each of the 5 clusters from C.

To determine the relationship between age-associated epigenomic changes and monocyte subpopulations, we combined differential regions from CM and NCM and clustered them based on the ATAC intensity. We identified 5 clusters: old CM specific, young CM specific, young CM shared with NCM, young NCM specific, and old NCM specific (**Figure 2C**). We observe few regions that are accessible, let alone differential in both subpopulations. Next, we ran HOMER to identify enriched motifs in each of the 5 clusters (**Figure 2D**; **Supp Figure 2B**). As with LOLA, we find AP-1 motifs in cluster 1 regions increased in aging CM. In addition, CTCF motif is enriched in both cluster 3 (decreased in aging CM) and cluster 5 (increased in aging CM). We also newly find that KLF and CEBP motifs were enriched in regions decreased in aging CM and NCM, respectively (clusters 2/3 and 4). To narrow down which KLF family member is driving this signal, we compared the expression of the associated gene and downstream targets. KLF4 was both expressed highly in CM and its downstream genes were downregulated in aging. As confirmation, we used JASPAR to validate the binding profiles of specific TFs in the aging signature (**Supp Figure 2C**). We confirmed that the CTCF binding profile was associated with regions decreased in aging CM and increased in aging NCM (p=0.002, p<1×10^−10^). On the other hand, we found that regions with CEBPA demonstrated the opposite trend in the aging signature (i.e. increased in aging CM p=4.9×10^−9^, decreased in aging NCM p= 4.7×10^−5^) which explains the discrepancy between LOLA and HOMER results. Among AP-1 family members, we chose ATF3 which has been previously associated with chromatin reprogramming in aging^45^: we found that it was enriched in regions with increased accessibility in both monocyte subpopulations (p=5.5×10^−10^ (CM), 2.7×10^−8^ (NCM)). Unfortunately, JASPAR did not have a binding profile for KLF4. Finally, we assessed the binding profile of PU.1 as a general TF in myeloid cells and found that, as expected, it was not enriched in the aging signature (*data not shown*). Our findings support the role of multiple factors, possibly with opposing activity, in CM and NCM suggesting that their aging signature is driven by diverse intrinsic and extrinsic signals.

### The aging signature of monocytes is dependent on TNF signalling

Prior work from our group demonstrates that TNF has a significant impact on aging monocyte development and function^37, 46^. In our current study, we included monocytes from TNF-KO mice raised in parallel with wild-type (WT) mice to determine the impact of TNF on transcriptional regulation of aging CM and NCM. By comparing expression across all age-associated genes in the absence of TNF, we observed that not only did aging TNF-KO monocytes exhibit expression closer to young WT but also the change in expression between young and old TNF-KO monocytes was decreased (**Figure 3A**). Similarly, when we set a cut-off based on whether gene expression was closer to young or old WT (see Methods), we found that the majority of sites were TNF-dependent (CM=55.6%, NCM =70.5%; **Supp Figure 3A**). When we performed GO analysis as in Figure 1 and compared the results, we found that the majority of processes previously associated with the aging signature were enriched in the TNF-dependent subsets (**Figure 3B**). A possibly exception is the metabolism-associated processed that were upregulated in aging CM. We note that although the TNF-related pathways (such as *regulation of tumor necrosis factor production)* were only significant in NCM, individual genes that appeared to be increased in both subpopulations were reduced in TNF-KO mice (**Supp Figure 3B**). Overall, monocytes from mice lacking TNF signalling demonstrated a widespread reduction in aging signature on the transcriptional level.

**Figure 3.**
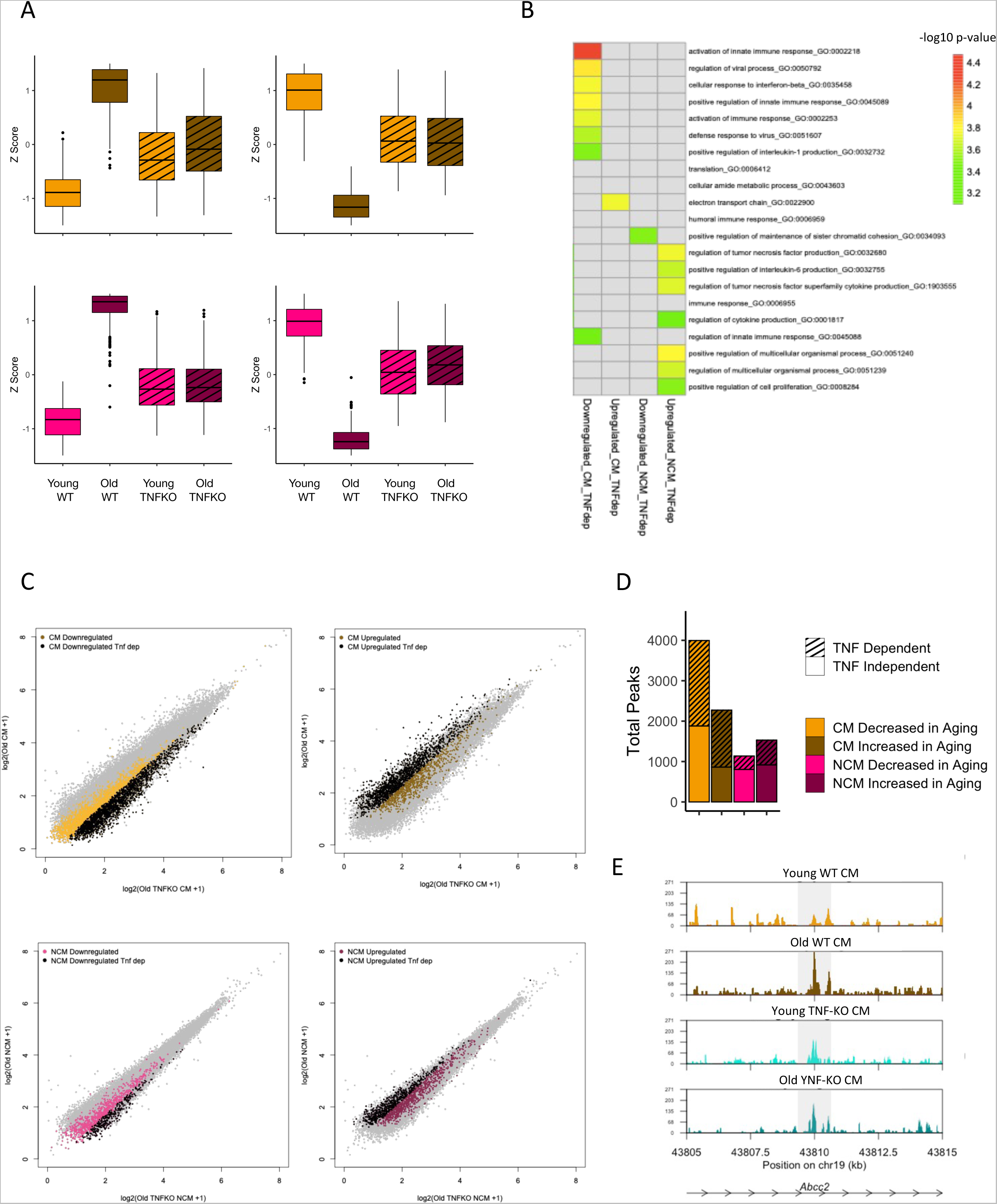
CM and NCM exhibit TNF-dependent aging signatures. A. Boxplots show Z-score transformed average expression values for genes that are increased (left) and decreased (right) in aging from Figure 1A across wild-type (WT) and TNFKO young and old CM (top) and NCM (bottom). B. Heatmap of gene ontology (GO) analysis results on the subset of TNF-dependent aging DEGs from Figure 1. C. Scatter plots of the average ATAC signal in log2 space for Old WT and Old TNF KO CM (top) and NCM (bottom). Regions that were increased (right) or decreased (left) in chromatin accessibility for each cell type (Figure 2B) are colored and shown in black if they are also TNF-dependent. D. The number of TNF dependent and independent regions in the aging signature as shown in (C). E. Genome tracks of chromatin accessibility at the locus near *Abcc2* in Young and Old WT and TNF-KO in CM.

Next, we analyzed the chromatin accessibility of monocytes from the TNF-KO mice. We found that both old TNF-KO CM and NCM exhibited distinct shifts in the accessibility of regions in the aging signature (**Figure 3C**). As with gene expression, we annotated peaks based on TNF dependency: in this case: NCM regions exhibited a much lower percentage of TNF-dependent regions than CM (35.5% vs. 56.3%; **Figure 3D**). We then looked at TNF dependency by cluster from Figure 2, but we did not observe any trends associated with regions across monocyte subpopulations (**Supp Figure 3C**). We also assessed the ATF3 binding profile specifically since it was linked to age-associated increases in both CM and NCM and found that regions with ATF3 sites were likely to resemble their young chromatin state in old TNF-KO mice (**Supp Figure 3D**). Some examples include the regions near *Abc22* and *Lmbr1* in CM and *Relt* and *Klf6* in NCM (**Figure 3E**; **Supp Figure 3E**). Therefore, TNF signalling is a critical element of the aging environment leading to monocyte dysregulation.

### TNF Over-expression Partially Mimics Age-Associated Changes to Epigenomic Landscape

In order to determine whether TNF alone can cause the observed changes in chromatin accessibility in the absence of the aging environment, we performed ATAC-seq on CM and NCM isolated from the bone marrow of transgenic TNF mice (TNF-OE) which express high levels of human TNF (hTNF) which binds TNF receptor 1 (TNFR1). We focused specifically on regions in the aging signature and filtered out those with low signal in monocytes from both TNF-OE and controls (**Supp Figure 4A**). We found 298 and 213 regions that were increased in both aging and TNF-OE mice (**Figure 4A top right**) and 643 and 81 regions that decreased in both aging and TNF-OE mice (**Figure 4A bottom left**) in CM and NCM, respectively. When we repeated the LOLA analysis from Figure 2, we found that the majority of TFs that were previously identified to have enriched binding sites still had significant overlaps with the combined aging/TNF-OE signature (**Figure 4B**). Next, we calculated the fraction of each quadrant that was labelled as TNF dependent (**Figure 4C**): we would expect that regions with aligned aging and TNF-OE signature (top right; bottom left) would exhibit higher levels of TNF dependency than those with opposing signatures (bottom right; top left). However, this does not hold true for upregulated regions in CM or down-regulated regions in NCM. Moreover, we do not observe differences in magnitude when TNF dependent regions are compared with TNF independent regions (**Supp Figure 4B**). Lastly, we revisited TNF-dependent ATF3 binding sites from Figure 3 and found that they were also upregulated in TNF-OE monocytes (**Figure 4D**). Together with the TNF-KO results, we conclude that TNF drives a portion of age-associated changes in monocytes; however, TNF does not work alone.

**Figure 4.**
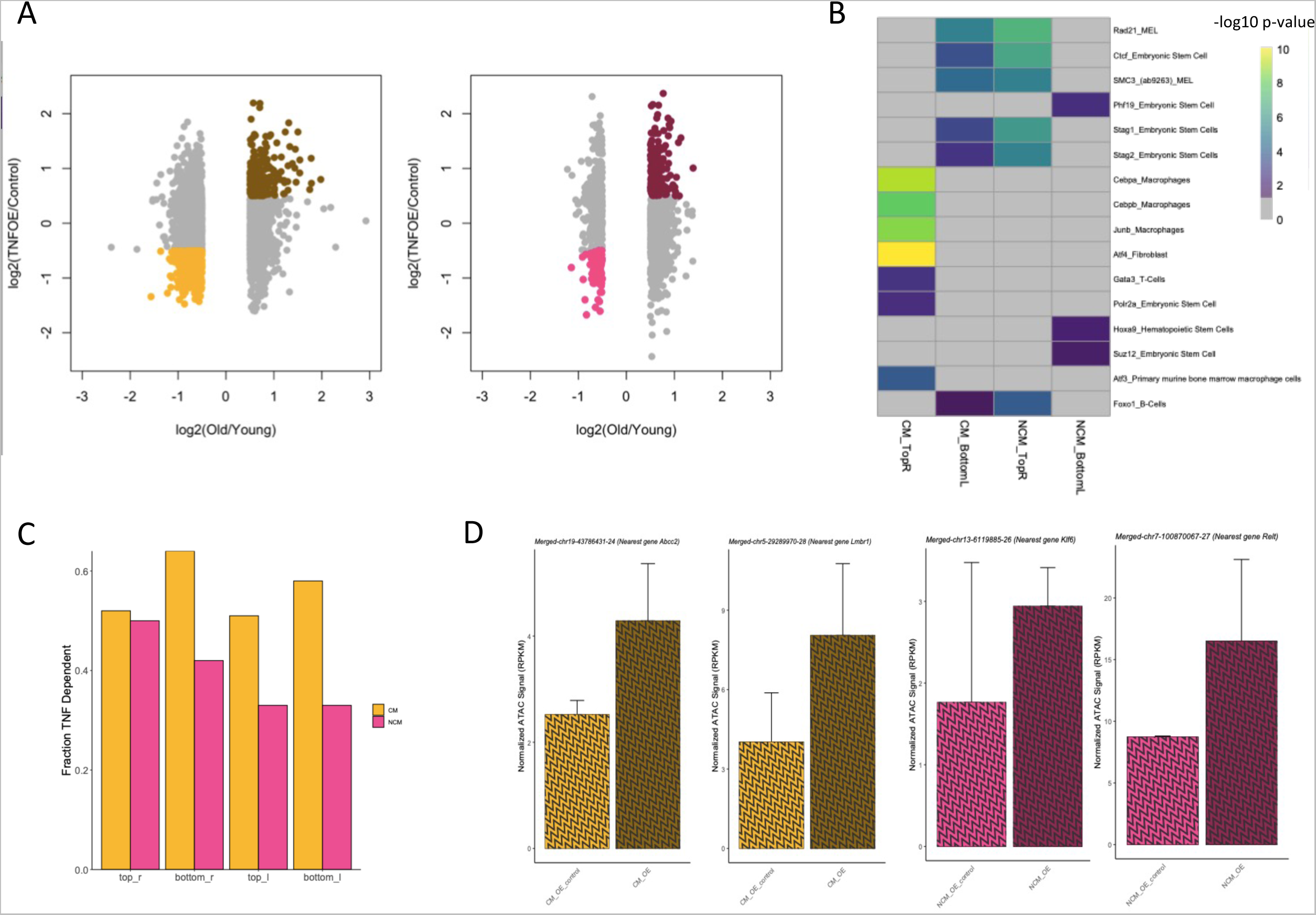
TNF overexpression (TNF-OE) signature overlaps the aging signature at a subset of chromatin regions. A. Scatter plots of log2 fold changes of ATAC-seq signal in aging signature regions between young and old mice (x axis) and TNF-OE and controls (y axis). Regions that are increased in aging and TNF-OE (top-right) and regions that are decreased in aging and TNF-OE (bottom left) are colored. B. Representative LOLA results for Top-Right and Bottom-Left regions in CMs and NCMs. C. The fraction of regions with aligned (top-right/bottom-left) or opposing (bottom-right/top-left) aging/TNF-OE signature from (A) that were annotated as TNF-dependent. D. Examples of ATF3 target regions with TNF-dependent increased accessibility in aging and TNF-OE.

## Discussion

Our study profiles monocytes from young and old mice to determine how monocyte transcriptional regulation is altered in aging. By using high-throughput genome-wide sequencing assays, we were able to characterize age-associated genes and regulatory regions in an unbiased approach. We determined that CM and NCM are impacted by aging in distinct manners. While TNF and other cytokine-related processes were upregulated in NCM, immune processes were surprisingly down-regulated in CM. It is possible that this non-intuitive trend represent a disruption of monocyte development in aging as these processes may be conflated with those involved healthy CM function. CM generally had a larger set of impacted genes and regions than NCM, which is consistent with our prior studies showing a greater increase in the numbers of Ly6chi monocytes ^37, 46^. These differences may reflect the fact that NCM are more terminal than CM. Since CM have the potential to further differentiate into circulating NCM and tissue macrophages, alterations to their transcriptional regulation may be passed on to have a systemic impact.

Our results indicate that similar TFs drive the aging signature in CM and NCM but they do not always act in the same direction. In particular, we found that binding sites for TFs involved in chromatin structure, such as CTCF and cohesion, were decreased in accessibility in CM and increased in NCM. CEBP binding and motifs exhibited the opposite trend. On the other hand, the AP-1 family TFs, including ATF3, were the key factors that were increased in both monocyte subpopulations. Importantly, none of these TFs bound the same sites across monocytes: they were enriched in unique regions of the CM and NCM aging signature. Thus, our approach of sorting the monocyte subpopulations is critical to understanding the impact of aging. It is possible that our reliance on external datasets not derived from monocytes and not equally available for all TFs obscured the binding trends. However, it was beyond the scope of this study to generate novel TF binding data, such as ChIP-seq, for individual factors.

TNF is involved in diverse processes from proliferation, differentiation, cell death, and inflammation. TNF is one of the key cytokines increased in expression in inflammaging, but its exact effect on cells is unknown. We showed an apparent reversal of the aging signature in mice lacking TNF signalling. However, TNF over-expression was not sufficient to recapitulate all the age-associated TNF-dependent changes we observed. This may be due to the fact that the TNF-OE model is based on hTNF, which, unlike murine TNF, does not bind TNFR2^47^. Another possibility is that TNF works in combination with other signals in the aging environment or cell-intrinsic factors to enact changes to monocytes. Our studies were unfortunately limited in their ability to distinguish between direct and downstream effects of increased TNF expression.

Aside from TNF, there are likely other factors that contribute to the aging signature of monocytes. This is supported by the significant number of TNF-independent genes and regions in our study. The fact that our aging signature included histone genes, chromatin-related GO processes, CTCF binding, and polycomb repressive complex activity suggests that aging is associated with large scale changes to chromatin organization in the monocyte nucleus. Prior studies have show a general dysregulation of the epigenome in aging including decreased histone occupancy, loss of heterochromatin regions, and differential DNA methylation ^9, 10, 48^. In addition, other groups described age-associated epigenomic changes that affect the differentiation of HSCs^18, 49^: these changes are likely propagated into the monocytes that develop from these HSCs. Moreover, aging HSCs are known to be myeloid biased and this in itself may drive defects in monocyte development in aging^21, 50^. While TNF and other signals in the environment may these epigenomic changes, it is also possible that they are cell-autonomous arising from intrinsic processes like DNA damage and accumulated mutations. In this case, the impact may be randomly distributed across the genome and not associated with specific processes or TF activity. It would also follow that these random aging effects would be heterogeneous within a given monocyte population. Thus, different approaches, such as DNA sequencing or single-cell RNA-seq, may be more appropriate to uncover these additional aging trends.

In summary, we demonstrated that aging monocytes exhibit altered transcriptional regulation that is, at least in part, TNF dependent. We also implicated specific biological processes and TFs in driving age-associated changes in the population-specific manner. These candidates may be further investigated in future studies to better elucidate upstream signals and downstream effects. Similar approaches could also be used to determine whether the aging signature is conserved in humans. Since monocytes are key to healthy immune function and response to challenge, understanding how they are dysregulated is critical to improving health in aging. Our study paves the way for future investigation of potential targets to ameliorate the effects of inflammaging and age-related diseases.

## Acknowledgements

We thank Dr. Hiam Abdala Valencia, Director of the Next Generation Sequencing Facility for the Divisions of Rheumatology/Pulmonary and Critical Care for sequencing services; the Lurie Flow Cytometry center for access to sorters; and the lab of Richard Pope for providing TNF-OE mice.

## Materials and Methods

### Mice

Young (3-6 months) and aged (20-24 months) female wildtype (WT) and TNF knockout (TNF-KO) mice on C57BL/6J background were aged for this study in a specific pathogen-free environment at the McMaster University Central Animal Facility (Hamilton, ON, Canada) as previously^37^. Aged mice used and bred for this study followed protocols and guidelines of Animal Research Ethics Board (AREB) of McMaster University. TNF-OE samples described in this manuscript are from the hTNF-TG mouse line on C57BL/6 background^51^ and were bred and raised in specific pathogen-free conditions in the Center for Comparative Medicine at Northwestern University Feinberg School of Medicine (Chicago, IL, USA) under an institutional animal care and use committee (IACUC) approved protocol.

### Bone marrow monocyte isolation and flow cytometry

At both McMaster (aging) and Northwestern University (TNF-OE), spine isolation and bone marrow extraction was performed from mice that were euthanized through cardiac exsanguination. Spine (from brainstem to tail) was dissected out of each mouse and cut into three pieces. Each piece of spine was opened and spinal cord was removed. Remaining pieces of spine were crushed with mortal and pestle in 10mL of cold PBS, pipetted through a 40uM cell strainer, and repeated until cells were clear. Single cell suspensions were spun down for 5min at 1500 rpm and resuspended in MACS buffer. Monocytes (XXX per mice) were isolated and sorted using conjugated antibodies of CD45+, CD11b+, CD19+, CD3+, CD56+, LY6G+, LY6C+/- sorted marker on an LSRII flow cytometer (BD Biosciences).

### RNA-seq library preparation

At least 50,000 bone marrow monocytes per subpopulation were sorted (as in **Supp Figure 1A**) and suspended in 200uL of RLT+ lysis buffer and stored at −80 degrees Celsius. Samples collected at McMaster were shipped overnight on dry ice to Northwestern University (Chicago, IL). RNA was extracted using Qiagen RNeasy extraction reagents and protocol. RNA-seq library preparation was performed with the QuantSeq 3’ mRNA-seq kit. The cDNA libraries were sequenced on Illumina NextSeq 500 platform with an average read depth of 20 million reads per sample and a 75-cycle single-end high-output sequencing kit.

### RNA-seq Processing and Analysis

Raw BCL files were demultiplexed using bcl2fastq2 software into FastQ files. FastQC was subsequently done to look at sample quality after sequencing. Raw reads were trimmed using bbduk (https://github.com/BioInfoTools/BBMap/blob/master/sh/bbduk.sh) and then aligned to the mm10 mouse reference genome using STAR^52^. Aligned reads were quantified and counted with HTseq to obtain a table of raw gene expression counts^53^. Count files were then normalized using CPM (counts per million). A final gene expression table was independently generated for CM and NCM by filtering out genes with a mean expression below 7.0 CPM in all experimental groups. Raw counts for these genes were used as input for the DEseq package^54^ to perform pairwise differential gene analysis. DEGs in aging were defined as those genes with p-value < 0.5 between young and aged mice. The lfcShrink() function set to type=”normal” was implemented to estimate log-fold change for comparisons and the output was plotted as a volcano plot using the R package ggplot. For Figure 3, we defined TNF dependency based on similarity of the Old TNF-KO expression to the Young WT (i.e. genes were considered TNF-dependent if |(Old TNF-KO) - (Old WT)| > |(OldTNF-KO) - (Young WT)|). Go Analysis was performed on genes within the aging signature data compared with classical or nonclassical background gene set using the program GOrilla^55^.

### ATAC-seq library preparation

ATAC-seq protocol was adapted from Buenrostro et al^56^ as described previously^57^ and initial steps were performed at McMaster (aging) and Northwestern (TNF-OE). Briefly, each sample consisting of approximately 50000 cells were spun down at 500xg, 4C, 20min in 0.2ml tubes, resuspended in 25ul of cold lysis buffer (10mM Tris-HCl pH 7.4, 10mM MgCl_2_, 0.1-0.5% IGEPAL CA-630), followed by another spin at 500_x_g, 4C, 20min. Cells were resuspended on ice with Transposase reaction (Buffer2x, Tn5, dH20 from NextEra) and then cleaned up by adding 5ul of clean up buffer (900mM NaCl, 30mM EDTA), 5% SDS and Proteinase K for 30min at 40C. All preparation was done on ice. Following the transposase reaction step, cells were frozen at −20C and for the aging experiment were shipped overnight on dry ice to the Winter Laboratory at Northwestern. ATAC-seq library preparation was completed at Northwestern University. Thawed cells were then cleaned up using a 2X AM Pure XP SPRI cleanup (Beckman Coulter A63881) with 100uL of 70% EtOH on Magnet. Supernatant were resuspended in 22ul of elution buffer (10mM Tris-HCl pH 8.0). I7 adapter index, I5 adapter index, Nextera PCR mastermix, PCR Primer cocktail, and template DNA was added to perform PCR (Polymerase chain reaction). PCR condition was run as 72C for 3min, 98C for 30sec, 98C for 10sec, 63C for 30sec, 72C for 3min, and 10C to store. Steps 3-5 ran for 9 cycles. Second SPRI size cut off was done on amplified PCR product with .65X AM Pure XP SPRI beads with same wash and magnetization as above. A second PCR amplification step was done following another 2X SPRI clean up. DNA libraries concentrations were measured by Qubit Fluorometer (Thermo Fisher) and the Agilent 4200 TapeStation. Libraries were sequenced using the Nextera Sequencing kit targeting 20 million paired-end reads per sample.

### ATAC-seq Chromatin Data Processing and Analysis

Reads from ATAC-seq datasets were aligned to mm10 mouse reference genome with Bowtie 2.0 with the following parameters: --very-fast -x-p 6 -1. ATAC-seq peaks were called with HOMER package *makeTagDirectory* followed by the *findPeak*s function^58^. *findPeaks* was run on the following parameters: -center -minDist 1000 -F 0 -L 2 -C 0 -size 500. 110,522 called peaks were merged using the *mergePeaks* function in HOMER. Peak counts were assigned to genomic regions using featureCounts^59^ and then normalized using FPKM (Fragment Reads Per Kilobase Million). To ensure comparable normalization and comparison across all samples (WT, TNFKO, and TNFOE), a quantile normalization was performed using the following steps: 1) determine the rank of each column, 2) sort the input table from lowest to highest, 3) calculate the mean of each row, 4) substitute the means in the ranked table. The *annotatePeaks.pl* function from the HOMER suite of tools was used to perform peak annotation. Only peaks that were present in at least two replicates of any group were retained. Correlation and PCA analyses were done using the *cor()* and *prcomp(dataTable, center = TRUE, scale. = TRUE)* R functions respectively. PCA plots were generated using *ggplot().* The threshold for the aging signature between young and old peaks was |log2(foldchange)| = 0.5. To generate the heatmaps, the aging signature table was passed through a row-min and row-max rescaling using the following function. This rescaled table was used to determine the 5 clusters in figure 2 with the *kmeans()* function in R. The aging signature was then plotted according to these clusters with pheatmap. TNF dependency for the chromatin accessibility of regions was defined analogously to gene expression (i.e TNF dependent means |(Old TNF-KO) - (Old WT)| > |(OldTNF-KO) -(Young WT)|). For Figure 4 analysis, TNF-OE regions that overlapped the aging signature were further filtered for signal in TNF-OE and control samples (Figure 4A, minimum mean density in log2 space CM = 2; NCM = 1.5).

### Genome Tracks

To generate the genome tracks, a custom R script was written using GenomicRanges R package. Total reads mapping to each chromosome and RPKM normalization was performed for each chromosome length. This standardized normalization across all regions of a chromosome and facilitated consistent retrieval of genomic regions which were then plotted in the form of genome tracks.

### Overlap with Binding Profiles

LOLA^60^ was used to determine the overlap of the aging (Figure 2) and joint TNF-OE (Figure 4) signature with experimental binding sites compared with background of filtered peaks in CMs and NCMs. LOLA output p-values were merged across comparisons and plotted using pheatmap in R. Unbiased motif enrichment analyses in Figure 2 were done using the HOMER suite of tools^58^. The *findMotifsGenome.pl* function was implemented with -*size given* and using the 48771 filtered and annotated peaks as the genomic background. Bed files of the binding profiles of TFs of interest were downloaded from the JASPAR database (https://jaspar.elixir.no/)^61^. Each transcription factor binding profile was filtered to a p-value cutoff of 10^−9^. These were entered as input with ATAC peaks from the aging signature to determine overlap using bedtools intersect with at least a minimum of 1bp overlap. The DoRothEA^62^ package (https://saezlab.github.io/dorothea/articles/dorothea.html) was used to obtain a list of targets for each TF in the KLF family. Then, the mean expression difference in aging was calculated to represent the effect on downstream genes.

---

**Figure 1 Supplemental.**
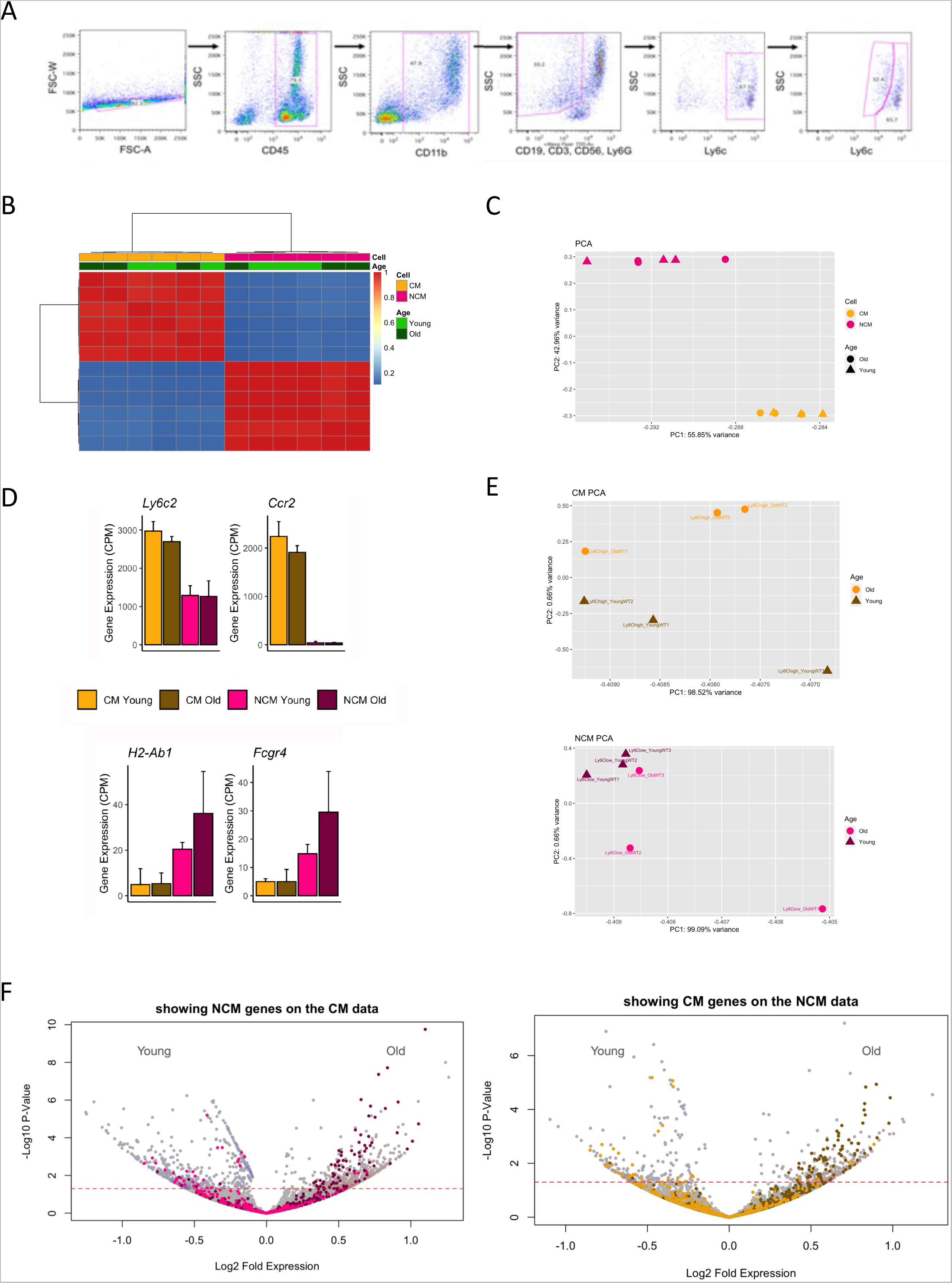
A. Flow cytometry panel used to isolate Ly6chi (CM) and Ly6c low (NCM) monocytes from bone marrow B. Adjacency matrix based on pairwise Pearson correlation of gene expression between RNA-seq replicates. C. PCA plot of gene expression colored by cell type (CM and NCM). D. Representative genes with differential expression between CM and NCM. E. PCA plot of gene expression colored by age (Old and Young) in CM (top) and NCM (bottom). F. Volcano plots of gene expression changes between Young and Old in CM overlayed with NCM genes (left) and in NCM overlayed with CM genes (right).

**Figure 2 Supplemental.**
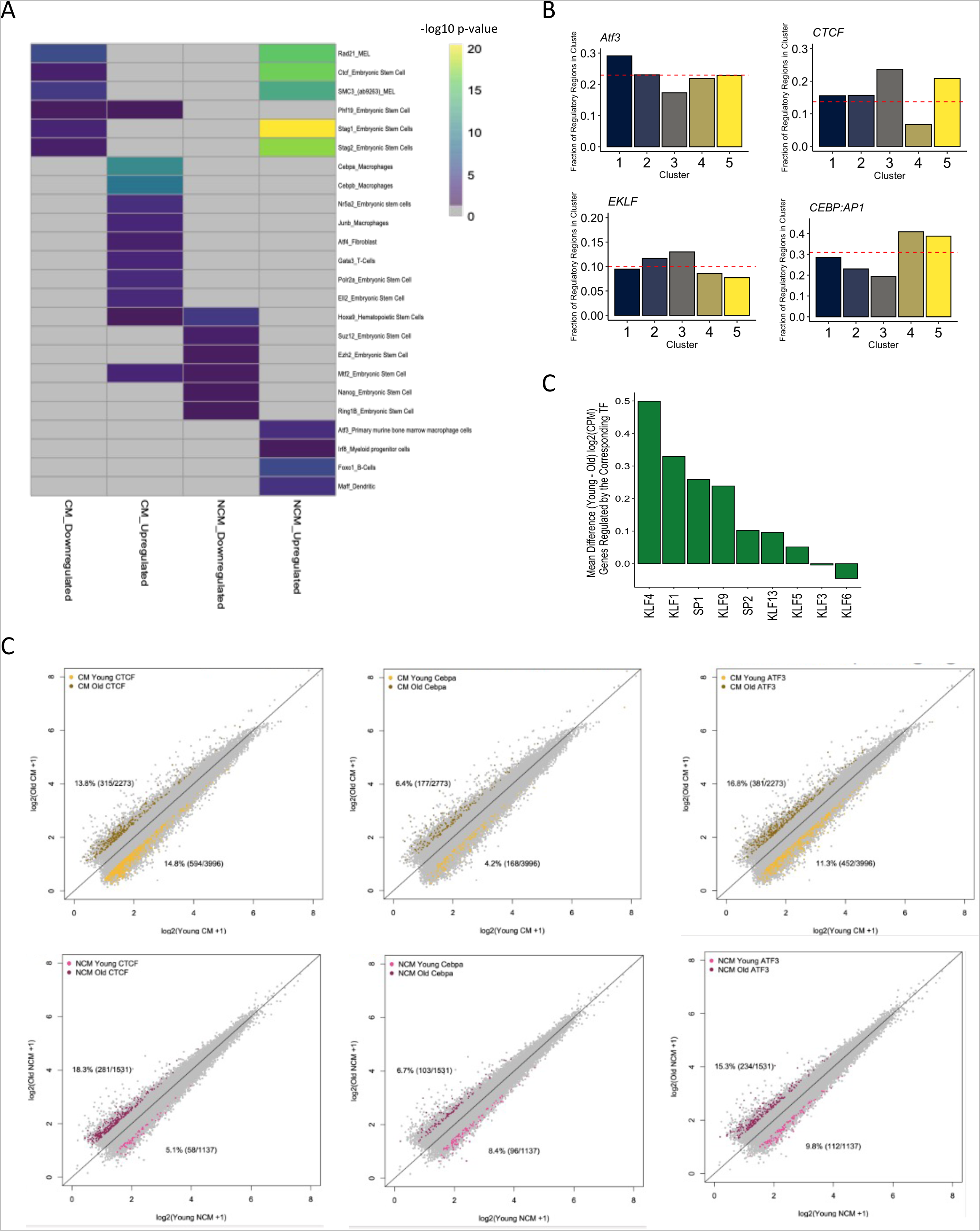
A. Representative LOLA results for aging signature of chromatin accessibility including CM down, CM up, NCM down, and NCM up. B. Summary of HOMER results across Figure 2C clusters for specific TFs. C. The mean difference in expression between young and old CM of downstream genes regulated by the given TF. C. Scatter plots of the average ATAC-seq signal in log2 space for young and old CM (top) and NCM (bottom) overlayed with aging signature regions overlapping JASPAR binding profiles for CTCF (left), CEBPA (middle), ATF3 (right).

**Figure 3 Supplemental.**
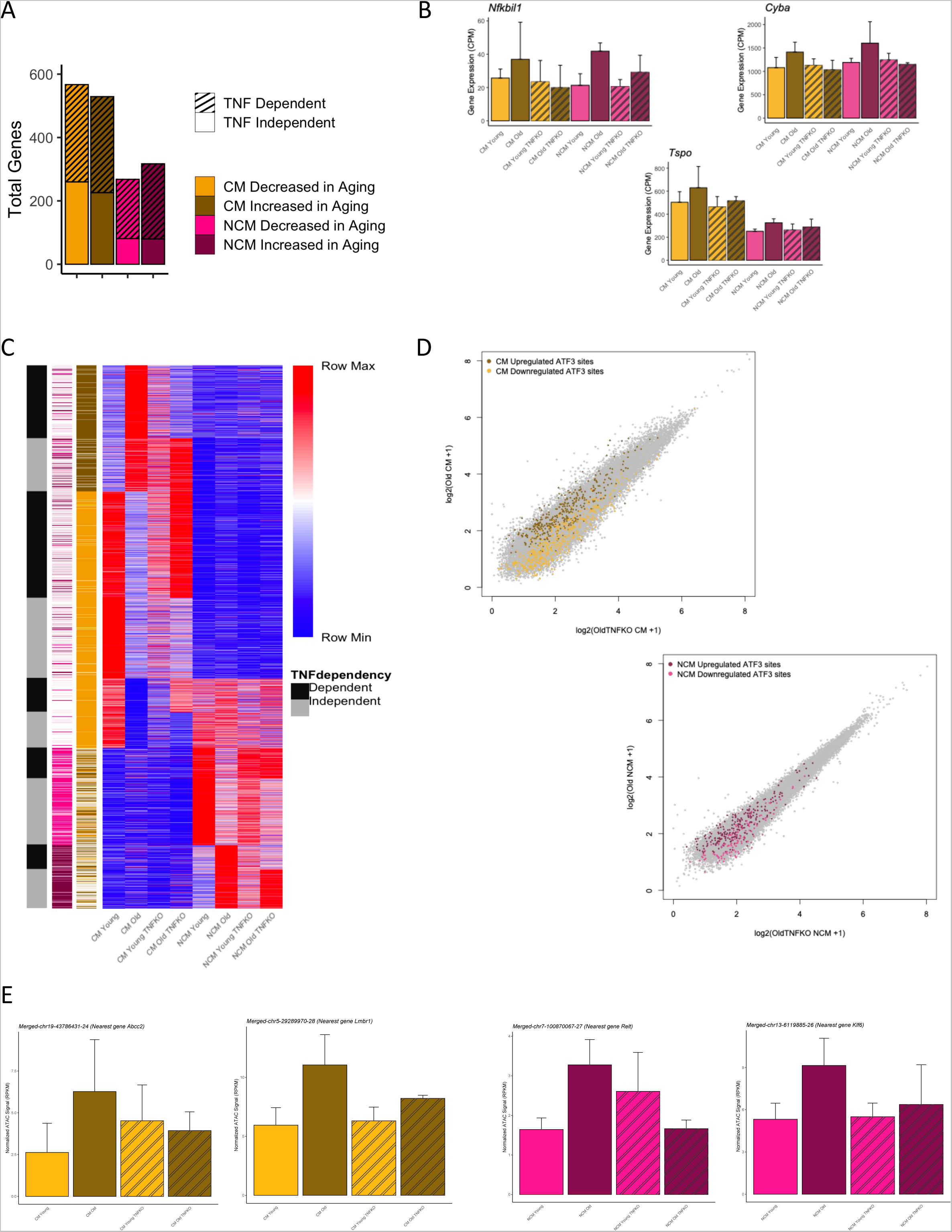
A. The number of TNF dependent and independent DEGs in aging CM and NCM corresponding to the analysis in Figure 3B. B. Representative examples of DEGs from the GO process *regulation of tumor necrosis factor production* that exhibit a TNF-dependent increase in aging NCM. C. Heatmap showing mean ATAC signal of regions across young and old WT and TNF-KO CM and NCM from clusters in Figure 2C split by annotation of TNF dependency. D. Scatter plots of the average ATAC signal in log2 space for Old WT and Old TNF KO CM (top) and NCM (bottom) overlayed with aging signature regions (increased in aging CM=brown, NCM=maroon; decreased in aging CM = yellow, NCM = pink) overlapping the JASPAR binding profile of ATF3. E. Mean normalized ATAC signal across young and old wild-type and TNF-KO monocytes in representative regions increased in aging CM (left) and NCM (right) that overlap the ATF3 binding profile.

**Figure 4 Supplemental.**
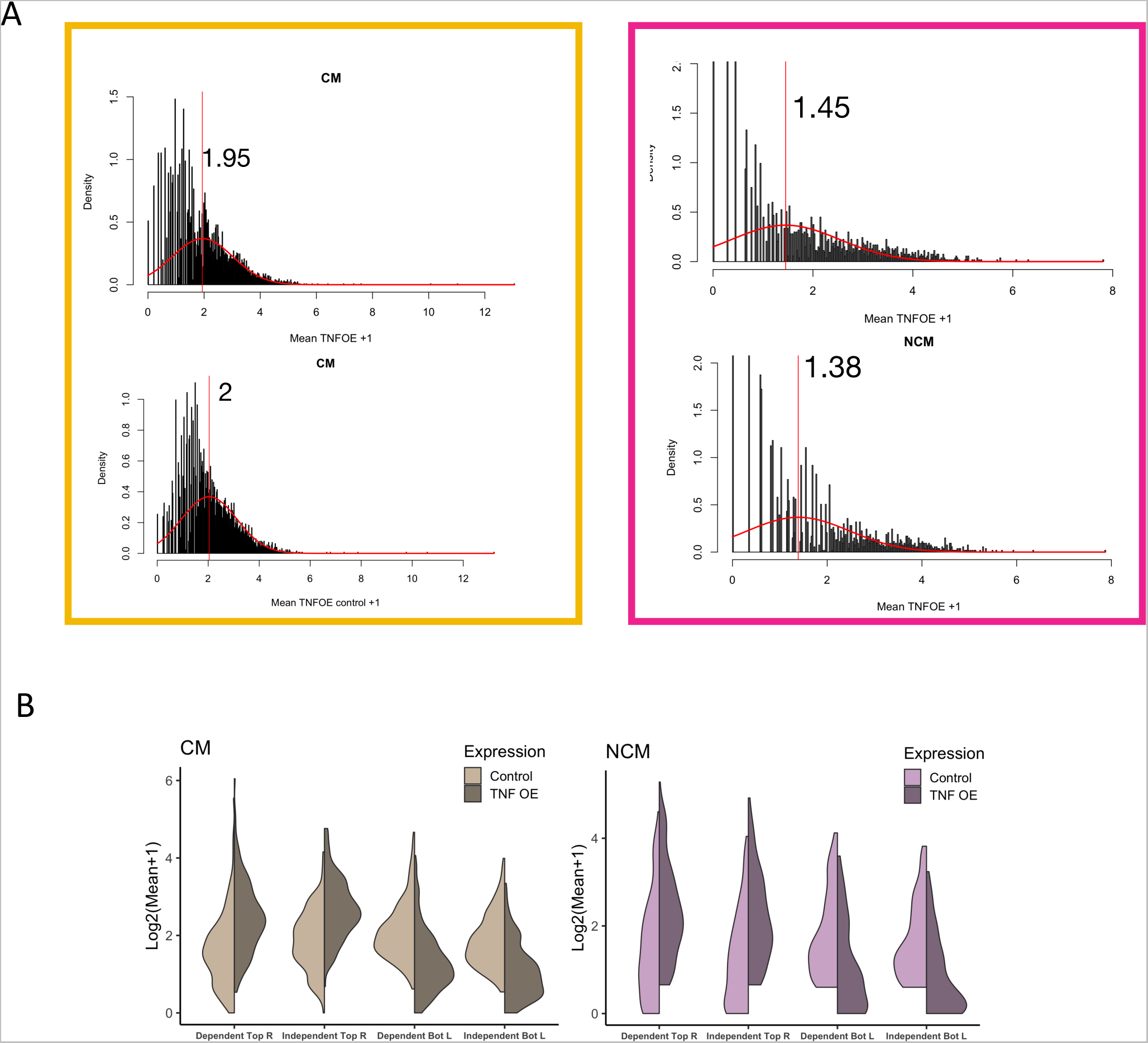
A. Distribution of ATAC-seq in CM (left) and NCM (right) from TNF-OE (top) and Control (bottom) mice. The red curve indicates the modelled distribution of regions with true signal with the red line at the calculated mean value. B. Violin plots of ATAC-seq signal in regions with aligned aging/TNF-OE signature split by TNF-dependence.

